# Graph-based learning and read-out of nanopore translocation event signals

**DOI:** 10.64898/2026.04.29.721625

**Authors:** Mayukh Kansari, Tobias Ensslen, Jan C. Behrends, Maria Fyta

## Abstract

Nanopores enable single-molecule analysis by measuring current signals through nanoscale pores in either biological or solid-state membranes. Accurate detection of analyte fingerprints within the pore environment is essential for reading-out the analyte type. We develop a framework for robust and label-free detection of the molecular nanopore events using a graph representation of the measured signals. To this end, we build a graph-based two-stage workflow based on a convolutional and graph neural networks that first perform a fast screening of the nanopore events, followed by a deep validation of these. The learned model can thus efficiently and in an unsupervised manner select possible molecular signatures (the current blockades) in the full signal, denoise, validate, reconstruct these, and predict the morphology of unseen molecular events. We could show that the learned model can efficiently predict the correct event morphology for the same analyte within a 2.4-fold range of transmembrane voltage values not included in the training. The developed graph-based workflow is modular, generalizable, and provided that it is trained on a huge amount of different nanopore experiments has the potential to become a blueprint model for nanopore read-out. Such a read-out model would be able to identify subtle differences in molecules like proteins, as well as their conformational or folding states. The proposed framework is developed using experimental signals from DNA translocation through an aerolysin pore and demonstrates a unified approach linking unsupervised feature learning to raw-signal inference for single-molecule sensing.

## I. INTRODUCTION

Nanopore technologies utilize nanometer-sized holes in solid-state or biological membranes to electrophoretically guide charged biomolecules such as DNA, RNA, or proteins [1–11]. The resulting passage of these analytes through the pore creates transient modulations (“current blockades”) in the ionic current measured across the pore. These time-dependent current signals serve as sensitive measurements of the analyte’s physical and chemical attributes, such as size, conformation, and sequence, establishing nanopore sensing as a robust platform for single-molecule detection and sequencing [12, 13]. The experimental current traces are though complex, can include high noise levels and baseline fluctuations, which render simple methods, such as traditional thresholding or segmentation, inadequate for robust detection of the blockades or molecular translocation events though the pore. Accordingly, a rigid obstacle in nanopore measurements is the reliable detection of these events, which is a critical step towards full read-out of the molecular identity. To this end, an error-free read-out demands sophisticated read-out algorithms capable of signal preprocessing, noise mitigation, and accurate event detection and classification. These tasks can be handled by Machine Learning (ML) algorithms, such as Deep or (DNNs) [14, 15], Convolutional Neural Networks (CNNs) [16–18] or Hidden Markov Models (HMMs) [19, 20].

In nanopore read-out algorithms, detecting single events enables the extraction of key features such as amplitude, duration, and multi-level substructure from noisy ionic current traces [21, 22], which correspond to single-molecule features. First event detection methods, such as MiniAnalysis and EasyNanopore [23, 24], relied on manually setting global current thresholds. These were often unreliable due to substantial noise and baseline drift inherent in high-bandwidth recordings [25, 26]. In order to improve robustness, adaptive methods were developed, utilizing techniques such as the Cumulative-Sums (CUSUM) algorithm for rapid, automated abrupt change detection as in OpenNanopore [27–29] and subsequent level fitting within detected events [29]. Other schemes, like the Transalyzer [28] involved the iterative baseline correction to overcome moving-average weaknesses by iteratively removing detected events and refining the mean and standard deviation [28, 30]. AutoNanopore [25, 31], a highly adaptive statistical outlier identification, uses Interquartile Ranges (IQR) and quantiles instead of mean and standard deviation to accurately locate events amidst fluctuating baselines [25, 30]. Further refinements include the Dynamic Correction Method (DCM), which employs sliding windows to dynamically compute local mean and variance, generating a nonlinear threshold curve to resist interference from baseline fluctuations and dense pulse signals [26]. Advanced methods can precisely locate event boundaries by monitoring the sign change of the signal slope rather than the baseline crossing [31] or by fitting a Fourier series to the event and locating the event extrema using second derivatives as in the Second-Order-Differential-Based Calibration [30, 32].

Novel read-out schemes are based on complex deep learning (DL) architectures, such as CNNs and Bi-Path Networks [14, 33, 34]. These claim a parameter-free event detection and classification and are often trained on extensive synthetic datasets [21, 35]. Other specialized ML approaches include the use of 1D CNN based on ResNet for classifying event-containing regions [35, 36] or a hybrid pipeline combining Incremental Learning (IL) to adapt base-calling for modification-rich sequences [37]. For base-calling raw signals directly, models like Ravvent combine raw signal and event features using Bidirectional long short-term memory (BiLSTM) encoders with an attention mechanism [38], demonstrating the move towards joint processing for enhanced accuracy.

Generally, these and other nanopore read-out and event detection schemes read directly the time series data from the experiments. Contrary to those, in this work, we explore and leverage the capabilities of Graph Neural Networks (GNNs). These extend deep learning principles to handle data residing on complex and irregularly structured graphs [39–42]. The core operation defining a GNN layer is mathematically realized as a graph convolution operation through the use of a graph filter [39, 41, 42]. This process requires the conversion of the graph signal from the vertex domain into the graph spectral domain using the Graph Fourier Transform (GFT), effectively decomposing the data onto a graph frequency basis [39– 41]. Through these spectral coefficients, GNNs can implement functions analogous to classical filtering, enhancing the denoising and smoothing of signals on the graph [39–41]. Graph Convolutional Networks (GCNs) perform local neighborhood averaging, which serves as a linear approximation of local spectral convolution [41–44]. In the biological context, GNNs could enhance the signal-to-noise (SNR) of fMRI data [39, 40], perform feature extraction from brain activity signals [39], and perform prediction in the area of health monitoring [42].

Building upon the demonstrated success of feature-engineered ML approaches [39–41], we propose employing GNNs as a novel methodological framework for analyzing nanopore ionic blockade currents. GNNs are uniquely adept at modeling complex relationships and structural dependencies inherent within time-series data, offering a powerful capability for robust pattern recognition. By leveraging the inherent strengths of GNNs, we aim to develop a more efficient and resilient method to accurately detect translocation events from the ionic current blockades obtained during nanopore sequencing experiments.

## II. METHODOLOGY

This work introduces a graph-based representation for event detection, contributing a new direction in the development of robust and scalable nanopore read-out algorithms. We introduce a novel methodology to handle the complexity of nanopore signals by integrating advanced neural network architectures with feature learning. We couple a GNN to an autoencoder framework applied directly to the detected translocation events. This combined architecture is designed to implicitly extract highly meaningful features by encoding the complex temporal and structural characteristics of the ionic blockades into a reduced-dimensionality representation known as the latent space. This learned feature space provides a robust environment for unsupervised analysis, enabling us to efficiently cluster the events and distinguish distinct molecular signatures or conformations. Following this successful feature extraction and verification step, the same trained GNN parameters are leveraged to operate directly on raw ionic data, performing the critical task of event detection in a unified and highly accurate manner, circumventing the limitations associated with traditional thresholding or segmentation methods. For demonstrating the applicability and efficiency of the developed GNN scheme we use nanopore data of DNA oligomers through a biological nanopore. We next present the main methodological details of the algorithms and technical details used in our workflow.

### A. Dataset Preparation and Annotation

Electrophysiological recordings were acquired from ssDNA made of 5 adenosines, (5A) translocating through aerolysin nanopores. The nanopore experiments were conducted in a 4 M KCl solution under transmembrane bias of 100 mV. All experimental details can be found elsewhere [45]. The recorded ionic current signals are stored in proprietary dat format and exported as axon binary format (abf) files. Raw current traces consisted of baseline open-pore segments interspersed with transient current blockades due to DNA passage. Translocation events were detected in traces using a threshold-based algorithm developed by the experimentalists, producing a curated reference set of 7,624 positive events with durations between 0.5–100 ms.

In order to generate negative samples for binary classification, regions of stable open-pore current (absolute deviation less than 0.5 *σ*, lasting at least 30 ms) were extracted from the baseline current, with windows randomly sampled to match the positive event duration distribution (0.5–30 ms), yielding 7,624 negative samples after exclusion of any true events. All waveforms were resampled to a standard length of 20,000 points via linear interpolation, and normalized using *z*-score scaling:

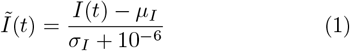

where *Ĩ* (*t*) is the normalized current, *I*(*t*) the measured current, *µ*_*I*_ and *σ*_*I*_ are mean and standard deviation. The latter two parameters account for amplitude variability and baseline drift. The resulting dataset was partitioned into training/validation/testing sets with a ratio of 70/20/10 using stratified sampling to maintain class balance.

### B. Two-Stage Event Detection and Representation

In this work, we develop a two-step workflow for our read-out algorithm. This consists of a hierarchical analysis based on an initial CNN screener for high-throughput candidate detection (stage 1), followed by a Graph Autoencoder (GAE) for event validation, morphological analysis, and unsupervised clustering (stage 2). This process is followed to learn and reconstruct the raw nanopore data in order to provide an automatized and highly accurate detection of events in these data. Briefly, the raw current waveform is segmented into overlapping windows via a sliding window approach. The windows, that is the candidate events, are screened at stage 1 and passed to stage 2 and are reconstructed. The original and reconstructed data are compared in order to validate the candidate events. The two-stage workflow is schematically provided in Fig. 1, while its key details are discussed in the following.

**FIG. 1.**
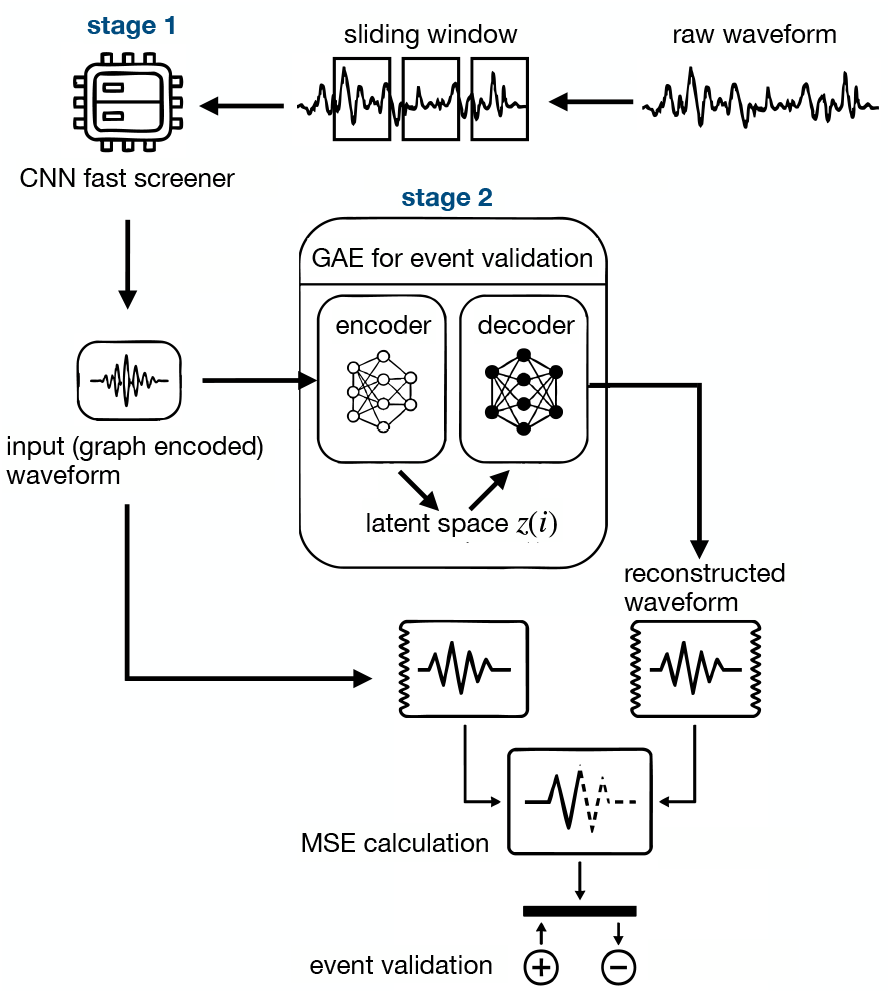
Schematic overview of the two-stage inference pipeline for nanopore translocation event detection and validation. A raw current waveform is segmented into overlapping windows via a sliding window approach. Each window is passed through Stage 1 and the CNN fast screener for the event candidate classification. The classified waveforms are fed into Stage 2 and the trained Graph Autoencoder, including the CNN encoder for compressing the normalized waveform, the GCN that refines each embedding through message passing through the *k*-nn, and a CNN decoder that reconstructs the waveform from the refined latent code. The MSE is used to compare the original input to the reconstructed waveform in order to validate the candidate as as true (+) or false (−) translocation events (+), respectively (see text for details).

#### 1. Stage 1: Convolutional fast screener of event candidates

The screener is a compact binary classifier composed of two one-dimensional convolutional layers (kernel sizes 5, 3 with 16, 32 filters, respectively), each followed by a ReLU activation function and a 4-fold max-pooling. A final fully connected layer employing a sigmoid activation function yields the output probability. This lightweight CNN outputs a binary event probability classifying a window as an event candidate or not. This architecture is trained using positive and negative events for scanning the events faster. The decision threshold was conservatively set (*τ* = 0.45) to prioritize high recall (*>* 95%). For inference, the network was applied in sliding windows of 20,000 samples with a 2,500 sample stride. Overlapping predictions were merged using an intersection-over-union criterion (*τ*_IoU_ = 0.4). Boundary refinement was performed by extending candidate windows based on current deviations greater than 30 pA. To facilitate rapid screening of extended electrophysiological recordings without reliance on manual thresholding, we designed a compact one-dimensional convolutional neural network (1D-CNN) to serve as a binary event detector. This architecture was intentionally optimized for throughput and high recall, operating under the paradigm that downstream, graph-based validation would remediate any false positives retained during primary screening.

Given any normalized waveform input **x** ∈ ℝ^20,000^, the screener performs the following mathematical transformations:

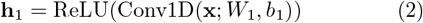

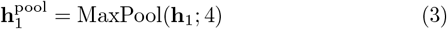

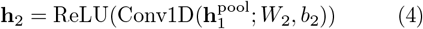

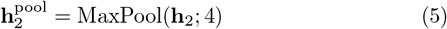

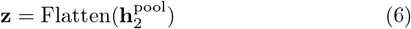

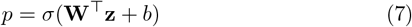

with kernel size 5 (at layer 1) and 3 (at layer 2), zero-adding used to maintain length. *σ*(·) is the sigmoid ctivation. **h**_**i**_, *W*_*i*_ represent the output and weights of each layer *i, b* the bias, **h**_**i**_^pool^ is the output of the *i*-layer, while *z* and *p* define the input and output of he latent space. The resulting flattened feature vector encodes broad morphological properties and global devitions, providing discriminative power for binary classification at minimal computational cost.

The CNN screener was trained on the balanced set of 15, 248 segments (50% true positive and 50% false negative events), using the binary cross-entropy (BCE) loss for *N* samples:

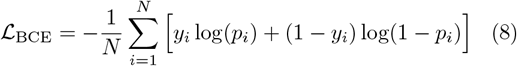

with the Adam optimizer and *η* = 0.005 for 550 epochs and early stopping based on validation loss. *y*_*i*_ and *p*_*i*_ define the output and respective probability. To prioritize event recall, the validation metrics (MSE) were calculated at a deliberately low operating threshold (*τ* = 0.45). For extended signals, the model was deployed with a sliding window (*w* = 20, 000, stride 2, 500 and 75% overlap) to ensure robust detection of events at window boundaries. Overlapping positive windows were merged using an intersection-over-union criterion (*τ*_IoU_ = 0.4). Event boundaries were extended to encompass all samples deviating by at least 30 pA from baseline.

The role of the CNN screener in the two-stage cascade process is to act as a gating function, reducing the noise from the dataset by up to 95% enabling the graph model to concentrate on difficult, morphologically ambiguous cases. Its adaptive, data-driven decision boundary is more flexible than static amplitude thresholds, allowing generalization across experimental variations, provided the training distribution is appropriately representative of future use.

#### 2. Stage 2: Graph Autoencoder for event validation

The filtered/classified event candidates from stage 1 are further encoded in stage 2 as nodes in graphs. The edges in this graph representation map the similarity of the event candidates. Stage 2 is supported by a Graph Autoencoder, which is made of a CNN encoder that compresses the normalized waveform, a GCN that refines each embedding, and a CNN decoder that reconstructs the waveform from the refined latent space. In the graph representation, the node features comprise the CNN-generated, 128-dimensional embeddings of the normalized waveform through the latent vector *z*(*i*) of the latent space. The GCN refines each embedding through message passing over a fully connected *k*-nearest neighbor similarity graph (*k*-NN with *k* = 5 the number of neighbors) and a CNN decoder that reconstructs the waveform from the refined latent space. The graph topology was constructed via the cosine similarity, was dynamically updated every five epochs, thus capturing relationships between morphologically similar events. The autoencoder is made of a two-layer graph convolutional encoder, followed by a CNN decoder employing transposed convolution and bi-linear interpolation. This reconstructs the original waveform from the 128-dimensional latent representations. The model was trained using Adam optimizer with *η* = 0.001, using a combined loss function penalizing both waveform reconstruction error (mean squared error) and graph structure loss via link prediction (positive and negative edge sampling). An 80/20 training/validation edge split was used for unsupervised validation. At the end of stage 2, the original input waveform and the reconstructed waveform are compared using as a metric the mean-squared-error (MSE), which performs the final validation. The GAE framework consists of four interconnected modules operating in sequence as sketched in Figure 2:

**FIG. 2.**
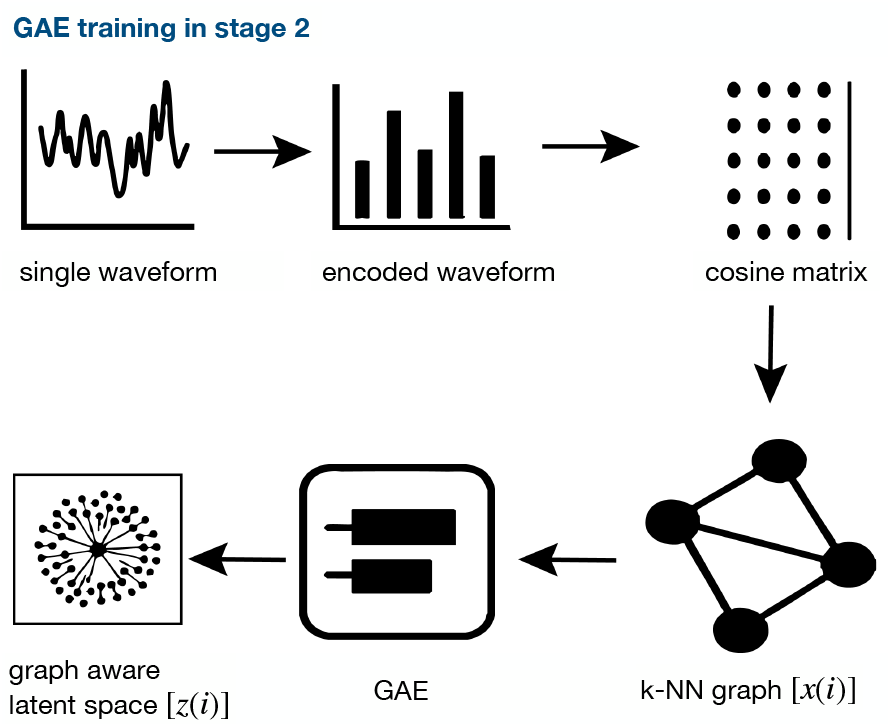
Schematic of the GAE training workflow for the learning graph-aware latent representations of translocation events. Each waveform (top left) is passed through a CNN encoder. The pairwise cosine similarities of these waveforms are computed across all encoded events to produce a similarity (cosine) matrix used to construct a *k*-NN graph with *x*(*i*) nodes is constructed. The graph is then processed by the GAE leading to a graph-aware latent space *z*(*i*) (bottom left) (see text for details).

1. **CNN feature encoder**: Initial feature extraction for each event candidate that compresses it into a 128-dimensional feature vector yielding the encoded waveform.
2. **Dynamic** *k***-NN graph constructor**: Similarity based graph topology based on the similarity (cosine) matrix extracting the five (*k* = 5) morphologically most similar neighbors to which each event node *x*(*i*) is connected.
3. **Graph convolutional encoder**: applies two layers of graph convolution to refine the embedding of each wavefront/event and output a graph-aware latent space. In this, each event occupies a unique point in a 128-dimensional manifold shaped by both its own waveform morphology and the structure of its surrounding event population.
4. **Convolutional decoder**: Waveform reconstruction from latent codes

#### 1. Convolutional feature encoder

Each normalized waveform 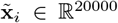 is first processed through a two-layer one-dimensional CNN to extract initial feature representations independent of graph structure. The encoder architecture consists of two layers. The first layer reduces the temporal dimension to 5,000 samples using 32 convolutional filters (kernel size 3, padding 1), a ReLU activation function and a 2-fold max-pooling. The second layer yields feature maps of length 2,500 using 64 convolutional filters (kernel size 3, padding 1), ReLU activation, followed by 2× max-pooling, yielding feature maps of length 2,500. The layers are fully connected, so that dense projection flattening the (64 × 2500) feature maps to a 128-dimensional embedding **h**_*i*_ ∈ ℝ^128^. With these characteristics, this CNN-based initialization captures local temporal patterns, such as rise times, peak structures, decay kinetics within single waveforms prior to graph-based refinement. The 128-dimensional bottleneck ensures computational tractability for subsequent graph operations, while retaining sufficient expressivity for morphological discrimination.

#### 2. Dynamic k-NN graph constructor

In order to explicitly capture the morphological similarity across the event dataset, a *k*-NN graph was constructed in the feature space 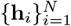 . For each event node *x*(*i*), edges were added to its *k* = 5 most similar neighboring events based on the cosine similarity:

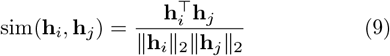

Note that the edges are bidirectional, yielding an undirected graph with adjacency matrix **A** ∈ {0, 1}^*N ×N*^ . The choice of *k* = 5 balanced the local neighborhood granularity with computational efficiency. Smaller *k* values risk fragmented components, while larger *k* dilute the morphological specificity. The graph topology is *dynamically updated* every 5 training epochs to reflect the evolving feature representations **h**_*i*_ as the CNN encoder weights are being optimized. This co-adaptation ensures that the graph structure remains consistent with the learned feature space throughout training, preventing stale topologies from constraining later-stage optimization.

## 3. Graph convolutional encoder

The CNN-derived features {**h**_*i*_} serve as initial node attributes for a two-layer graph convolutional network (GCN) for refining the wavefront representations by aggregating the information from the graph neighbors. The spectral graph convolution is then applied on each GCN layer as follows

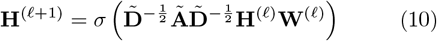

with **Ã** = **A** + **I** the adjacency matrix with added selfloops, 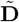 the degree matrix of **Ã**, with 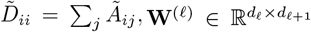 the learnable weight matrices, *σ* the ReLU activation function, and **H**^(𝓁)^ the node features at layer 𝓁, with **H**^(0)^ = {**h**_*i*_} . Both GCN layers include 128 hidden nodes, thus keeping the same dimensionality throughout the graph convolution stack. The symmetric normalization 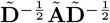 balances the message contributions from neighbors with varying degrees, preventing high-degree nodes from dominating the aggregation. The output of the second GCN layer results in the graph-aware latent representations **z**_*i*_ ∈ ℝ^128^. These include both the local waveform morphology (from the CNN encoder) and the neighborhood context (from the graph convolution). Accordingly, events with similar neighborhoods develop correlated latent codes, enforcing manifold smoothness conducive for clustering.

## 4. Convolutional decoder

In order to ensure that the latent space **z**_*i*_ retains sufficient information for an accurate waveform reconstruction, a CNN-based decoder projects the latent vectors back to the original 20,000-sample temporal domain. The decoder architecture thus mirrors inversely the encoder.

For the overall GAE optimization, the composite loss function is calculated:

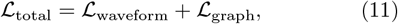

which balances the fidelity of the waveform fidelity and the preservation of the graph structure. The wavefront reconstruction loss (ℒ_waveform_) is defined as

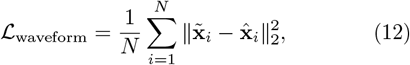

where 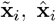 refer to the normalized input and the decoder output (reconstructed) waveforms. This mean squared error based error estimation encourages the latent space to preserve morphological information necessary for the waveform reconstruction. For low values of ℒ_waveform_, the primary supervision signal enforces accurate reconstruction of input waveforms from their latent representations.

The graph reconstruction loss (ℒ_graph_) is key for the prediction of the graph links. It ensures that the latent space retains the graph topology meaning that connected nodes (i.e. morphologically similar events) develop proximal latent information. Following standard GAE training protocols, the edge set ℰ was partitioned into positive edges ℰ^+^ (actual *k*-NN connections) and randomly sampled negative edges ℰ^−^ (non-connected node pairs). The graph reconstruction loss penalizes deviations from the expected similarity structure and implicitly maps an auxiliary link prediction:

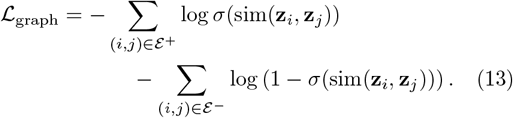

The similarity in the latent space is computed via the cosine distances:

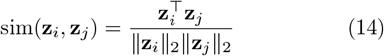

Accordingly, a high cosine similarity maps connected node pairs and low similarity disconnected pairs, embedding the graph topology into the geometry of the latent manifold.

For the validation of the candidate events at the end of the process in Figure 1 two fidelity metrics are used: the MSE for evaluating the amplitude preservation and the Pearson correlation for the shape similarity between input and reconstructed waveforms. Events with high reconstruction fidelity (r ≥ 0.7) and MSE ⩽ 2.9 are accepted as true translocation events (true positives), while poorly reconstructed candidates are rejected (false positives). These threshold values are obtained from the reconstructed events from the decoder during the training process, as most of the real events lied in this range. An additional restriction is set for the dwell time, that is the duration of each event. Accordingly, events in the range of 0.5 − 50*ms* are calssified as true events. The latent space generalization of the training set is quantified by computing the Euclidean distance between each inferred event and its nearest neighbor. At the same time, any events above *µ*_*d*_ + 2*σ*_*d*_ (training mean and standard deviation) are considered outliers. The performance of the developed workflow is assessed based on the held-out test set by Stage 1 recall, Stage 2 precision, and cluster consistency (fraction of identical cluster assignments for the nearest neighbors in the latent space).

## III. RESULTS AND DISCUSSION

### A. Features and event embeddings onto graphs

After the two-stage screening, 3,388 translocation events were retained from the initial candidates using a screening threshold of *τ* = 0.45. The threshold was set slightly below the neutral sigmoid boundary (*τ* = 0.5) to prioritize recall over precision as a default value, enabling broad noise filtering in Stage 1. Subsequent Stage 2 reconstruction-based validation effectively eliminated the residual false positives. In order to visualise the transformation of the normalized wavefronts into the 128-dimensional feature vector through the CNN encoding, we randomly choose two morphologically very distinct wavefronts and depict these in Figure 3(a). As evident from the feature representation in panel (b), the two events map onto two distinct activation patterns, differing in both magnitude and waveform. Apparently, events with varying morphological characteristics, single-level or multi-level blockades, etc. produce distinct activation patterns across the 128 feature dimensions, confirming that the encoder captures morphologically distinct features. Despite the 156:1 compression ratio (20,000 dimensions are reduced to 128), the embedding in the 128-dimensional space captures the essential degrees of freedom of the translocation signal, while discarding redundant temporal correlation.

**FIG. 3.**
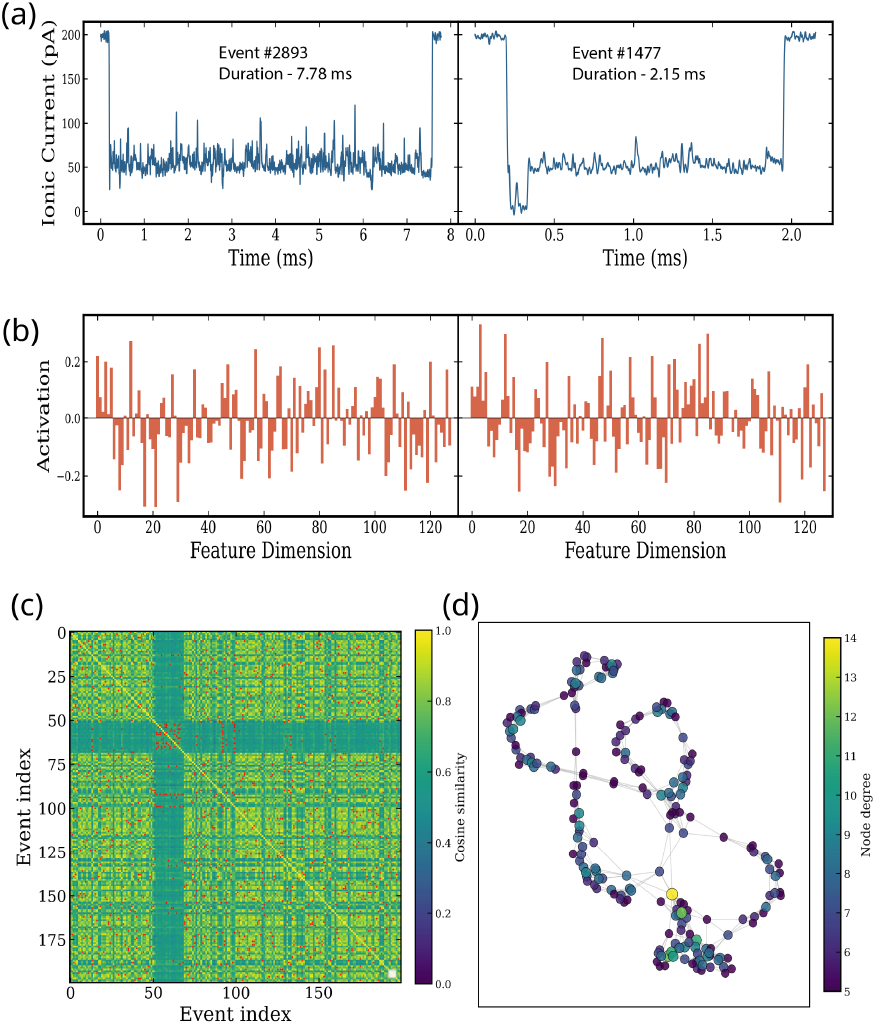
(a) Two representative and distinct translocation events: a long deep (50 pA) and a short double-level blockade. (b) The corresponding 128-dimensional CNN embedding vectors are displayed activation bar plots in the feature dimension. (c) The pairwise cosine similarity matrix for a representative subset of 200 events, with values ranging from 0.0 (dark green) to 1.0 (yellow) (see color bar). The block-diagonal structure reveals groups of mutually similar events. (d) Two-dimensional projection of the *k*-NN graph, with node colour encoding node degree (number of connections).

In order to visualize the morphologically distinct and similar events in the total signal the dynamic 5-NN graph was constructed from the CNN embeddings using the pairwise cosine similarity matrix for a representative subset of 200 events in Figure 3(c). The block-diagonal structure of this reveals groups of mutually similar events. Red dots mark the positions of the 5-NN, which concentrate within high-similarity blocks and avoid cross-block connections, demonstrating that the graph construction faithfully captures the local similarity structure of the embedding space. In panel (d), the projection of the 5-NN graph reveals a heterogeneous topology. Densely connected hub nodes (degree 12–14 depicted in yellow) reside in high-similarity regions where many events share similar waveform morphology, while peripheral nodes (degree 5–6 depicted in dark purple) represent morphologically distinct or rare events. This topological variation reflects the underlying diversity of the translocation event population and provides the structural scaffold for the subsequent GCN message-passing stage. The graph is rebuilt every 5 training epochs, allowing the connectivity to co-evolve with the learned representations. As the CNN encoder im-proves, similarity estimates become more accurate, the graph rewires accordingly, and the GCN operates on an increasingly faithful neighborhood structure.

### B. Event morphologies

The two-layer GCN refines the CNN embeddings into 128-dimensional latent vectors. A t–SNE projection (perplexity = 30) of the full latent space of 3, 388 events based on clustering with *k*-means (*k* = 4) is shown in Figure 4(a) revealing clear morphological similarities among events. Due to the nature of the homopolymer consisting of five-adenines, the clusters shown reveal subtle differences reflecting stochastic differences in translocation dynamics, such as fluctuations in the dwell time and current blockade depth, rather than fundamentally distinct ionic current signatures arising, which are expected from experiments including heterogeneous analytes. In the figure the four clusters are marked together with the number of events mapped in these. The cluster boundaries are well defined and most of the events are parts of the cluster structures with no significant outliers. Representative examples from each cluster are shown in Figure 4(b), including single- and multi-level, as well as highly noisy events. The first cluster (C1 in blue) includes short-to-intermediate duration events (2–5 ms) with clean, well-defined blockades and sharp baseline recovery. The second cluster (C2 in red) maps intermediate-duration events (3–10 ms) exhibiting deep blockades with variable sub-level complexity, noisy plateaus, and occasional multi-step transitions. The third cluster (C3 in pink) represents long-duration events (10–25 ms) characterized by extended, noisy current reductions with pronounced morphological complexity and gradual transitions. In the fourth cluster (C4 in cyan), the events reveal deepest blockade depths but short length compared to the other clusters, displaying a distinct amplitude profile that separates them from the rest of the populations in the latent space. These distinct group of events could point to low capture rates and incomplete translocation events.

**FIG. 4.**
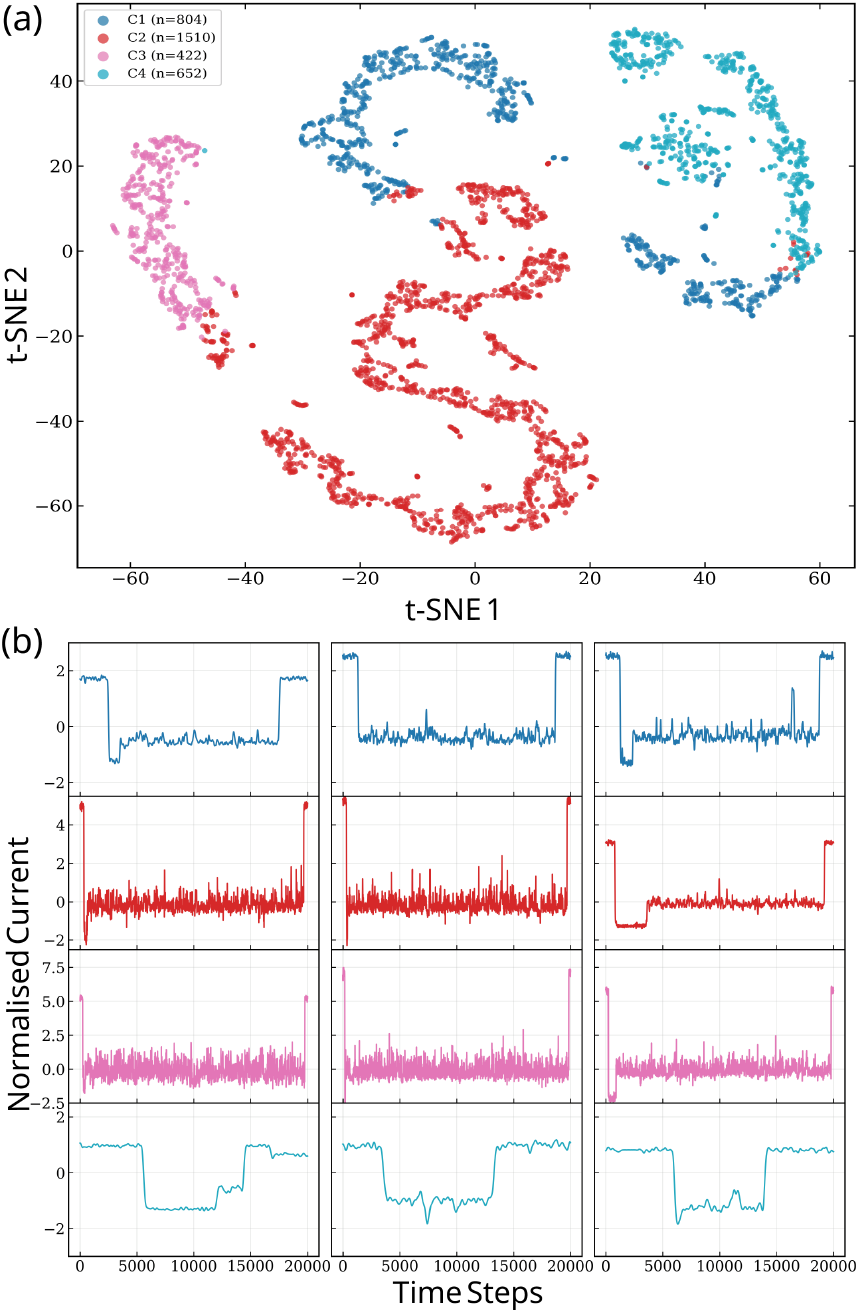
a) t–SNE projection of the 128-dimensional latent space with colored clusters denoted as C*i*, with *i* the cluster number. In parentheses, the respective number of events in the cluster are provided. (b) Three representative z-normalized waveforms per cluster are displayed matching the color coding in (a). For details see text.

In order to assess whether the latent space encodes physically meaningful properties, the clusters in the t–SNE map in Figure 4(a) are provided with respect to the two features event duration and blockade depth in Figure 5(a), essentially mapping the latent space. This refers to a non-linear dimensionality reduction scheme embedding high-dimensional data into a low-dimensional space revealing correlations. This representation reveals again the direct correlation of the clusters with long, short or intermediate events. The projection of the t–SNE2 component with respect to event duration in panel (b) reveals a smooth gradient orthogonal to the depth gradient, with short events occupying one end of the manifold and long events the other end. In the blockade depth case on the right of this panel, a smooth spatial gradient across the latent manifold with shallow events concentrated in the upper-left region is visible. A smooth gradient is visible across the manifold, with short events (*<* 5 ms) concentrated in one region and longer events (*>* 15 ms) occupying a distinct but continuously connected region. The respective distributions of the events are shown in Figure 5(c) for the duration and blockade depth. The different duration ranges discussed above are evident here, also revealing the most probable duration for each cluster with clusters C1 and C4 dominated by events with a median duration of 3–4 ms nad C3 revealing a median of 15 ms. The left panel shows that thress clusters (C1, C2, C3) exhibit comparable blockade depths (120–140 pA), while C4 displays a lower depth 30–50 pA. Note that in the the t–SNE projection in panel (b), the low-depth events occupy the top-left region of the manifold, spatially separated from the deeper blockades. The results indicate that the GAE learns the physical representation of the molecular translocation events mapping them into the 128-dimensional latend space efficiently and without supervision.

**FIG. 5.**
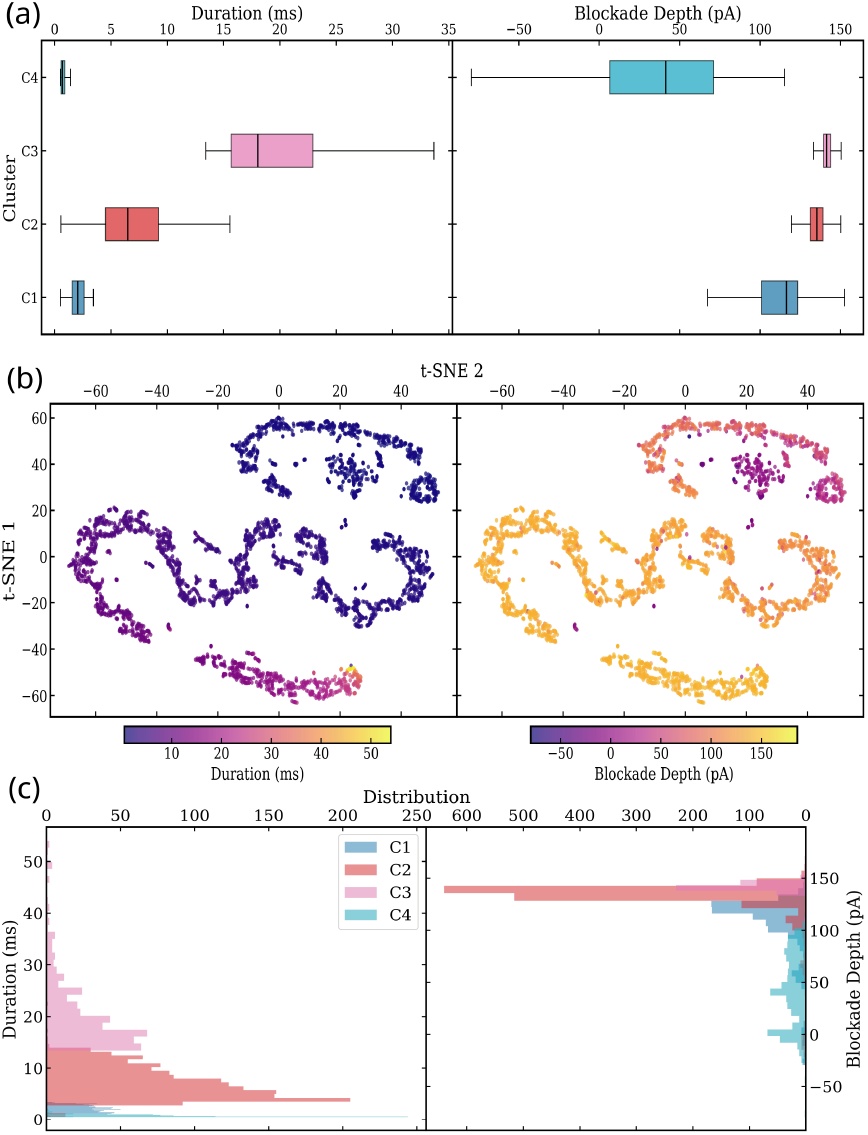
(a) Bar plots with respect to the cluster with the (left) event duration and (right) the blockade depth and the t–SNE2 component. The respective t–SNE2 correlation to these features is provided in (b), respectively. (c) The event distributions with respect to the duration (left) and the blockade depth (right).

### C. Event Reconstruction

The waveform decoder was next used for the reconstruction of the original events to serve as an additional assessment of whether the latent space indeed retains sufficient information for faithful signal recovery. Figure 6 compares six representative reconstructed events spanning the morphological diversity of the dataset to the original events. In all the main features (onset/offset of the event, blockade depth, baseline recovery, even low-frequency noise) and the overall morphology of the event are very well retained. This is quantitatively confirmed by the MSE and Pearson correlation coefficients (r) provided in the figure. For the selected events, the MSE values range from 0.0233 to 0.1557 and the correlation coefficients from r = 0.939 to r = 0.990. Implicitly, the reconstruction acts as an implicit low-pass filter on the signal. High-frequency stochastic noise present in the raw trace is absent from the reconstructed one. This behaviour is a direct consequence of the bottleneck architecture: the 128-dimensional latent code has insufficient capacity to encode random noise but sufficient capacity to encode the systematic signal component.

**FIG. 6.**
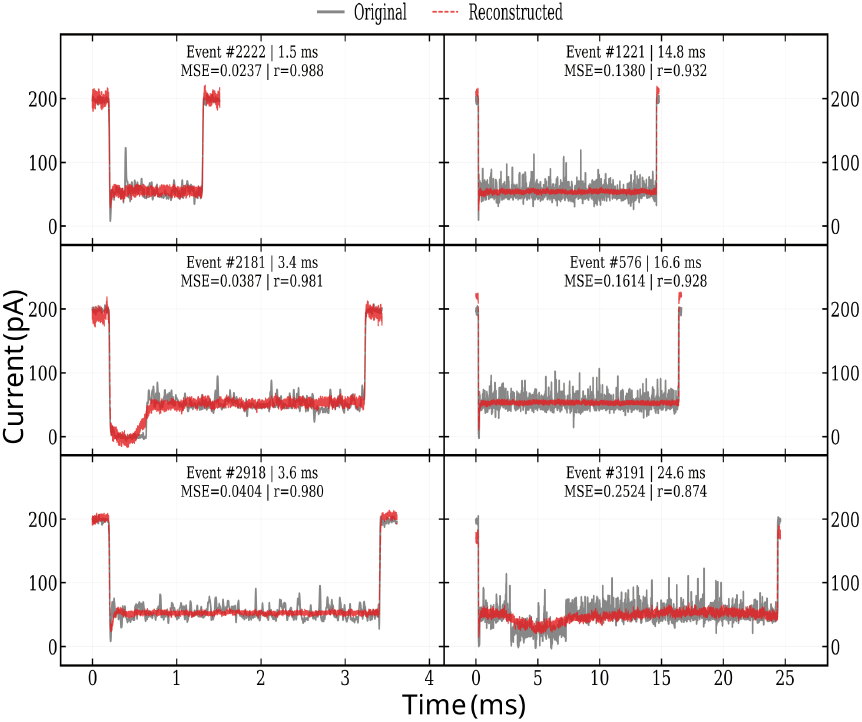
Six representative translocation events as reconstructed from the latent space (red) and compared to the raw normalized input (real) events. The MSE and Pearson coefficient r are provided, as well as the event number and duration.

Note that events of different morphologies are reconstructed with the same accuracy. Refer to the figure and compare short, single-level to the long, multi-level blockades with extended noisy plateaus. Their reconstruction was performed at the same accuracy level. In the case of noisy events (also depicted), the accuracy is reduced as the reconstruction does capture the underlying signal, but discards the stochastic component. The reconstruction efficiency of the latent space and the decoder ensures that during training the latent space retains the event waveform rather than collapsing into a degenerate representation. Later on, at the inference level, the quality of the reconstructed events provides an event-by-event validation score as detected events not well reconstructed (with *<* 0.7, MSE *>* 2.9) are classified by the Stage 1 screener as false positives and are excluded from the downstream analysis.

### D. Inference and event prediction

Once trained, the two-stage model was applied to the 100 mV recording used for training. Accordingly, the train model is tested. For this, the continuous raw signal, a 10-second segment of which having a 200 pA baseline is depicted in Figure 7(a). detected events highlighted in red. The Stage 1 CNN screener, operating in sliding-window mode (window = 20,000 samples, stride = 5,000), could successfully identify the dense population of translocation events. The detected events overlaid on the raw signal correspond to 93% of the expected events and confirms that the screener achieves high recall across the segment, capturing events of varying duration and depth while leaving baseline regions intact. The representative single events in panel (b) of the same figure are quite distinct morphologically and confirm the high quality inference on the training set. Specifically, events with clean, well-defined waveforms are reconstructed with high fidelity (r *>* 0.95, MSE *<* 0.05). Events with prolonged noisy plateaus or complex multi level ones show reduced but still informative reconstruction quality (r = 0.982-–0.988, MSE = 0.07—0.03). Note that for these lower-quality reconstructions, the decoder correctly captures the event envelope (onset, approximate depth, recovery), but not the high-frequency noise which is not kept in the latent space, as discussed also for the training process above.

**FIG. 7.**
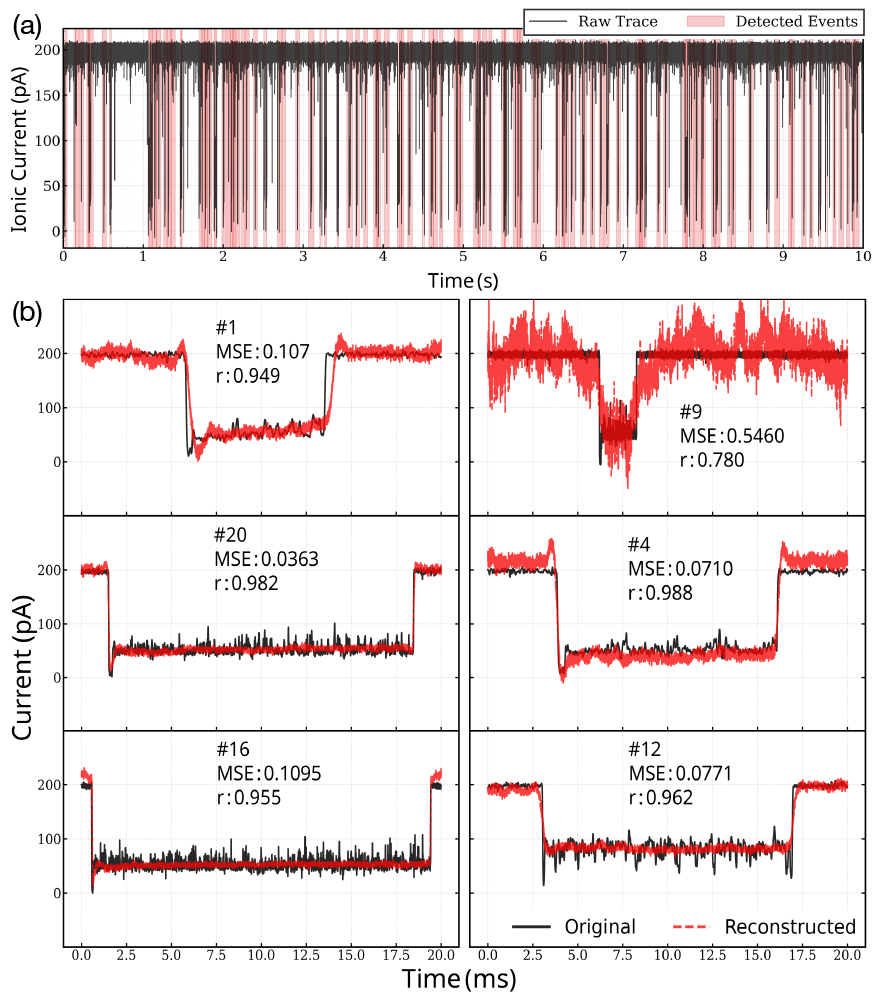
End-to-end inference on the 100 mV training recording. (a) 10 sec segment of the raw current trace (in black) and a 200 pA open-pore baseline with the detected translocation events highlighted in red. (b) Representative detected single events shown at full temporal resolution. The event number, the MSE and Pearson coefficient r are provided. The input/raw signals are provided in black and the reconstructed in red.

The inference pipeline is next generalized to new unseen voltage conditions. For this, the transferability of the developed workflow and the model trained on the 100 mV recordings is assessed. This model without further training is processing three different nanopore experiments with the same analyte and pore, but distinct applied voltages of 50 mV, 80 mV, and 120 mV. These conditions generate translocation events with different features than the trained ones at 100 mV. Representative detected and reconstructed events for these different conditions using the model trained at 100 mV are provided in Figure 8. It is clear, that the reconstruction accuracy remains high across all three voltages, with r in the range 0.908–0.988.

**FIG. 8.**
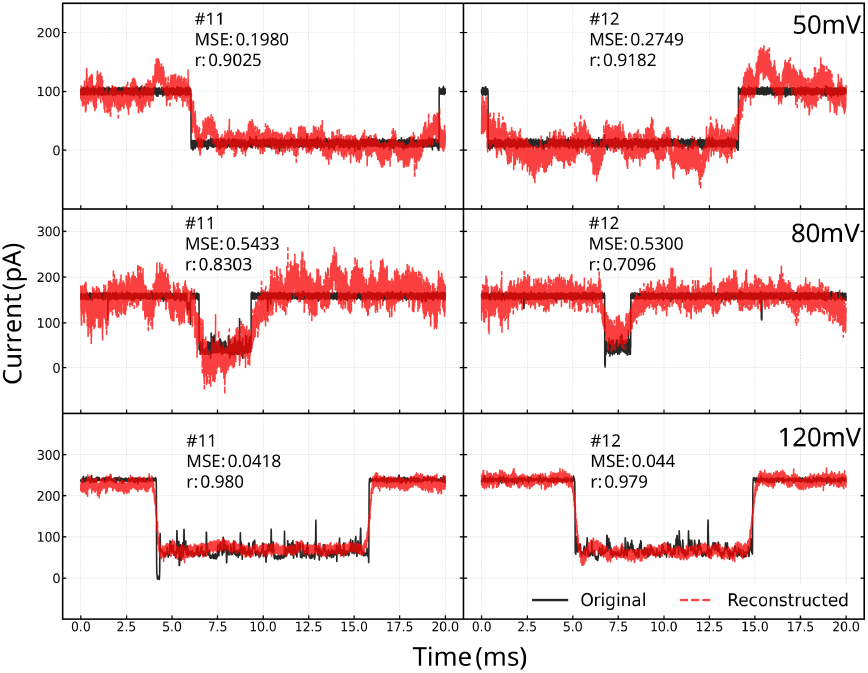
Representative events detected from the original signal (black) and as GAE-reconstructed (red) from 50 mV (top), 80 mV (middle), and 120 mV (bottom) recordings using the model trained for 100 mV recordings. The event number, the MSE and Pearson coefficient r are provided.

Specifically, at 120 mV, the events exhibit deep blockades and well-defined transitions and are reconstructed with high accuracy (refer to the accuracy metrics in the figure) capturing the multi-level sub-structure and the sharp onset edges. The higher driving force produces cleaner signals with higher SNR and the reconstruction quality reflects this accordingly. At 80 mV, the events are shallower and have an increased noise relative to the signal amplitude. Despite the model never having seen 80 mV data during training, it successfully detects events and produces accurate reconstructions (refer again to the accuracy metrics in the figure). The decoder correctly identifies the event waveform even when the blockade depth is substantially lower than the 100 mV training distribution. At 50 mV, the reconstruction quality is similar to that at 80 mV and is the most challenging as the low voltage is linked to shallow, slow, and noisy translocation events that differ most strongly from the 100 mV training data. Still, the model, effectively detects and reconstructs events with reduced blockade depths and the respective waveform morphologies.

The successful generalization across a 2.4-fold voltage range (50–120 mV) without retraining demonstrates that the graph autoencoder learns shape-based representations that are robust to changes in absolute signal amplitude, event morphology, and noise characteristics. The z-normalization applied during preprocessing, which removes absolute amplitude and standardizes waveform shape, ensures that the CNN encoder focuses only on the waveform morphology and not the voltage-dependent current scale, enabling transferability across experimental conditions. This eliminates the need to retrain the model for each voltage protocol.

## IV. CONCLUSIONS

Graph representations of molecular translocation events through nanopores were successfully processed in this work through neural network architectures in order to detect, learn, and reconstruct the experimental signals. The developed workflow is self-supervised and efficiently maps the molecular events onto a low dimensional latent space that retains the main morphology filtering out high frequency noise. The workflow is composed of four independent modules, the CNN encoder, the GCN, the waveform decoder, and the CNN screener, all connected through defined interfaces (128-dimensional vectors and edge index lists). In essence, the latent space encodes smooth gradients of the most important features, such as the event duration, blockade depth, and waveform morphology. The learning process is thus interpretable and strongly connects to the fact that the reconstruction loss forces the latent space to retain the single-event details. At the same time, the link prediction loss enforces geometric consistency in the latent space by requiring that graph-connected events, that is the morphologically similar events, remain proximal.

The separation of event detection into a fast screening and a deep validation stage resolves a fundamental throughput challenge in nanopore data analysis. As raw recordings can include a huge number of current data points, of which a small fraction corresponds to translocation events, running the GAE on every window of the raw signal would be computationally very demanding. The Stage 1 screener operates instead as a lightweight binary classifier with deliberately high recall. A very strong point of the graph-based two state process developed here is its successful generalization for the same analyte across a 2.4-fold range of applied transmembrane voltage (50–120 mV) without retraining. This voltage-independent performance is a direct consequence of the z-normalization preprocessing, so that the encoder only receives information on the event morphology and not the scale. We expect from these findings that the cluster-wise event detection should also be transferable to different analytes as the one trained.

In short, the graph-based two stage workflow developed in this work has the ability to encode subtle correlations of the events in nanopore recordings learning the similarity of these for the same analyte. The networks encode abstract morphology features retaining though the information on physically intuitive features of the events, such as the duration, depth, etc, which confirms previous supervised studies [16, 46]. This capacity is essential for detection of biomolecules and the post-translational modifications therein, as the differences of the molecular blocks are very subtle. The additional generizability to large voltage ranges would be applicable in probing voltage-dependent translocation kinetics, without the need of additional experiments. In a longer perspective, this would also enable the identification of protein folding states (folded, unfolded, partially folded) without requiring explicit definitions of those states. Another strong point of the workflow is its modularity, as each of the four components can be separately exchanged by another network or learning scheme. Note that the graph-based workflow involving a sliding-window screening followed by a GAE validation and characterization can be extended beyond nanopores to other experiment types measuring rare, transient events in a continuous noisy signal. Provided that the GAE component would be trained on a large and diverse set of nanopore data for various analytes, pore types and diameters, solvents, and tranmembrane voltage differences it could provide a blueprint foundation model for the nanopore community, analogous to the role of large language models in text processing and image generation.

## V. ACKNOWLEDGMENTS

The computing time provided in the NHR Center NHR4CES at RWTH Aachen University (project numbers p0021740) is greatly acknowledged. This is funded by the Federal Ministry of Education and Research, and the state governments participating on the basis of the resolutions of the GWK for national high performance computing at universities. This work is part the nanodiagBW consortium (project number 03ZU1208BI and 03ZU1208AJ) funded by the Federal Ministry for Research, Technology and Space (BMFTR) within the Clusters4Future initiative. Funding from the German Funding Agency (Deutsche Forschungsgemeinschaft) and DFG - Project number 508324943 is greatly acknowledged.

## CONFLICTS OF INTEREST

There are no conflicts to declare.

## VI. DATA AVAILABILITY

The data generated and analyzed in this work can be made available upon any reasonable request. Experimental data were recorded by T.E. in the laboratory of J.C.B. under Physiologisches Institut der Universität Freiburg. The Machine Learning workflow in this work is implemented in Python and available on GitHub

## Notes

### Competing Interest Statement

The authors have declared no competing interest.

## References

[1] Meng-Yin Li, Yi-Lun Ying, Jie Yu, Shao-Chuang Liu, Ya-Qian Wang, Shuang Li, and Yi-Tao Long. Revisiting the origin of nanopore current blockage for volume difference sensing at the atomic level. Jacs Au, 1(7):967–976, 2021.

[2] Arya Krishna, Neethu Puthumadathil, Dheeraj Kumar Sarkar, Bibhab Bandhu Majumdar, Devika Vikraman, Jagannath Mondal, and Kozhinjampara R Mahendran. Controlling nanopore dynamics via loop stapling and unstapling for tunable substrate transport. ACS nano, 2025.

[3] Kyloon Chuah, Yanfang Wu, SRC Vivekchand, Katharina Gaus, Peter J Reece, Adam P Micolich, and J Justin Gooding. Nanopore blockade sensors for ultrasensitive detection of proteins in complex biological samples. Nature communications, 10(1):2109, 2019.

[4] Thanh Hoang Phuong Doan, Jasper P Fried, Wenxian Tang, Daniel Everett Hagness, Ying Yang, Yanfang Wu, Richard D Tilley, and J Justin Gooding. Nanopore blockade sensors for quantitative analysis using an optical nanopore assay. Nano Letters, 24(21):6218–6224, 2024.

[5] Chenyu Wen, Shuangshuang Zeng, Zhen Zhang, Klas Hjort, Ralph Scheicher, and Shi-Li Zhang. On nanopore dna sequencing by signal and noise analysis of ionic current. Nanotechnology, 27(21):215502, 2016.

[6] Mazdak Afshar Bakshloo, John J Kasianowicz, Manuela Pastoriza-Gallego, Jérôme Mathé, Régis Daniel, Fabien Piguet, and Abdelghani Oukhaled. Nanopore-based protein identification. Journal of the American Chemical Society, 144(6):2716–2725, 2022.

[7] Gerhard Baaken, Ibrahim Halimeh, Laurent Bacri, Juan Pelta, Abdelghani Oukhaled, and Jan C Behrends. Highresolution size-discrimination of single nonionic synthetic polymers with a highly charged biological nanopore. Acs Nano, 9(6):6443–6449, 2015.

[8] John J Kasianowicz, Eric Brandin, Daniel Branton, and David W Deamer. Characterization of individual polynucleotide molecules using a membrane channel. Proceedings of the National Academy of Sciences, 93(24):13770–13773, 1996.

[9] Chan Cao, Nuria Cirauqui, Maria Jose Marcaida, Elena Buglakova, Alice Duperrex, Aleksandra Radenovic, and Matteo Dal Peraro. Single-molecule sensing of peptides and nucleic acids by engineered aerolysin nanopores. Nature communications, 10(1):4918, 2019.

[10] Fabien Piguet, Hadjer Ouldali, Manuela Pastoriza-Gallego, Philippe Manivet, Juan Pelta, and Abdelghani Oukhaled. Identification of single amino acid differences in uniformly charged homopolymeric peptides with aerolysin nanopore. Nature communications, 9(1):966, 2018.

[11] Hadjer Ouldali, Kumar Sarthak, Tobias Ensslen, Fabien Piguet, Philippe Manivet, Juan Pelta, Jan C Behrends, Aleksei Aksimentiev, and Abdelghani Oukhaled. Electrical recognition of the twenty proteinogenic amino acids using an aerolysin nanopore. Nature biotechnology, 38(2):176–181, 2020.

[12] Naren Das, N Mandal, Praveen Kumar Sekhar, and Chirasree RoyChaudhuri. Signal processing for single biomolecule identification using nanopores: a review. IEEE Sensors Journal, 21(11):12808–12820, 2020.

[13] Yanan Jiang and Wei Guo. Nanopore-based sensing and analysis: beyond the resistive-pulse method. Science Bulletin, 60(5):491–502, 2015.

[14] David Dematties, Chun Wen, Mario D Pérez, Dong Zhou, and Shi-Li Zhang. Deep learning of nanopore sensing signals using a bi-path network. ACS Nano, 15(8):13289–13301, 2021.

[15] Zi-Xuan Wei, Yi-Lun Ying, Meng-Yin Li, Jie Yang, Jia-Le Zhou, Hui-Feng Wang, Bing-Yong Yan, and Yi-Tao Long. Learning shapelets for improving single-molecule nanopore sensing. Analytical chemistry, 91(15):10033–10039, 2019.

[16] Ángel Díaz Carral, Magnus Ostertag, and Maria Fyta. Deep learning for nanopore ionic current blockades. The Journal of Chemical Physics, 154(4), 2021.

[17] Yuk Kei Wan, Christopher Hendra, Ploy N Pratanwanich, and Jonathan Göke. Beyond sequencing: ma-chine learning algorithms extract biology hidden in nanopore signal data. Trends in Genetics, 38(3):246–257, 2022.

[18] Hongxu Ding, Ioannis Anastopoulos, Andrew D Bailey IV, Joshua Stuart, and Benedict Paten. Towards inferring nanopore sequencing ionic currents from nucleotide chemical structures. Nature Communications, 12(1):6545, 2021.

[19] Jacob Schreiber and Kevin Karplus. Analysis of nanopore data using hidden markov models. Bioinformatics, 31(12):1897–1903, 2015.

[20] Shujie Zhang, Wei Chen, Laibo Song, Xiaohong Wang, Weilun Sun, Pengyun Song, Ghazala Ashraf, Bo Liu, and Yuan-Di Zhao. Recent advances in ionic current rectification based nanopore sensing: a mini-review. Sensors and Actuators Reports, 3:100042, 2021.

[21] Pratima Upretee, Jan Fostier, Wouter Botermans, Koen Martens, Sanjin Marion, and Nilesh Madhu. Detecting translocation of dna nanostructures through nanopores: First steps towards structural barcode readout. Proc. 31st Eur. Signal Process. Conf. (EUSIPCO), pages 1205–1209, 2023.

[22] Zachary Roelen, Kyle Briggs, and Vincent Tabard-Cossa. Analysis of nanopore data: classification strategies for an unbiased curation of single-molecule events from dna nanostructures. ACS sensors, 8(7):2809–2823, 2023.

[23] Jing Tu, Hao Meng, LinLin Wu, Guohao Xi, Jiye Fu, and Zuhong Lu. Easynanopore: A ready-to-use processing software for translocation events in nanopore translocation experiments. Langmuir, 37(33):10177–10182, 2021.

[24] Daniel Pedone, Markus Firnkes, and Ulrich Rant. Data analysis of translocation events in nanopore experiments. Analytical chemistry, 81(23):9689–9694, 2009.

[25] Zepeng Sun, Xinlong Liu, Wei Liu, Jiahui Li, Jing Yang, Fang Qiao, Jia Ma, Jingjie Sha, Jian Li, and Li-Qun Xu. Autonanopore: an automated adaptive and robust method to locate translocation events in solid-state nanopore current traces. ACS omega, 7(42):37103–37111, 2022.

[26] Guohao Xi, Jinmeng Su, Jie Ma, Lingzhi Wu, and Jing Tu. A robust signal processing program for nanopore signals using dynamic correction threshold with compatible baseline fluctuations. Analyst, 150(7):1386–1397, 2025.

[27] C Raillon, P Granjon, M Graf, L J Steinbocka, and A Radenovic. Fast and automatic processing of multilevel events in nanopore translocation experiments. Nanoscale, 4(16):4916–4924, 2012.

[28] Calin Plesa and Cees Dekker. Data analysis methods for solid-state nanopores. Nanotechnology, 26(8):084003, 2015.

[29] C. Raillon, P. Granjon, M. Graf, L. J. Steinbock, and A. Radenovic. Fast and automatic processing of multi-level events in nanopore translocation experiments. Nanoscale, 4:4916–4924, 2012.

[30] Pratima Upretee, Wouter Botermans, Koen Martens, Sanjin Marion, Jan Fostier, and Nilesh Madhu. Methods for single-biomolecule translocation event detection from nanopore current signal: A review. IEEE Sensors Journal, 2025.

[31] Xinlong Liu, Zepeng Sun, Wei Liu, Feng Qiao, Li Cui, Jing Yang, Jingjie Sha, Jian Li, and Li-Qun Xu. Multilevel translocation events analysis in solid-state nanopore current traces. In 2022 IEEE International Conference on Bioinformatics and Biomedicine (BIBM), pages 1648–1653. IEEE, 2022.

[32] Chenyu Wen, Dario Dematties, and Shi-Li Zhang. A guide to signal processing algorithms for nanopore sensors. ACS sensors, 6(10):3536–3555, 2021.

[33] Nuria Celik, Pavel Damborský, Raffaella Fiammengo, Kanchana Nalluru, Silke Hage, Jaroslav Katrlik, Mark De Smet, Eddy Peeters, Fien Vaneven, Wendy Van Roy, et al. Deep-channel uses deep neural networks to detect single-molecule events from patch-clamp data. Commun. Biol., 3(1):3, 2020.

[34] Motohiro Tsutsui, Takashi Takaai, Kenichi Yokota, Tomoji Kawai, and Takashi Washio. Deep learning-enhanced nanopore sensing of single-nanoparticle translocation dynamics. Small Methods, 5(7):2100191, 2021.

[35] Jaise Johnson, Chinmayi R Galigekere, and Manoj M Varma. A solid-state nanopore signal generator for training machine learning models. arXiv preprint arXiv:2504.05466, 2025.

[36] Julian Hoßbach, Samuel Tovey, Tobias Ensslen, Jan C Behrends, and Christian Holm. Peptide classification from statistical analysis of nanopore sensing experiments. The Journal of chemical physics, 162(8), 2025.

[37] Ziyuan Wang, Yinshan Fang, Ziyang Liu, Ning Hao, Hao Helen Zhang, Xiaoxiao Sun, Jianwen Que, and Hongxu Ding. Adapting nanopore sequencing basecalling models for modification detection via incremental learning and anomaly detection. Nature Communications, 15(1):7148, 2024.

[38] Adam Napieralski and Robert Nowak. Basecalling using joint raw and event nanopore data sequence-to-sequence processing. Sensors, 22(6):2275, 2022.

[39] Rui Li, Xin Yuan, Mohsen Radfar, Peter Marendy, Wei Ni, Terrence J O’Brien, and Pablo M Casillas-Espinosa. Graph signal processing, graph neural network and graph learning on biological data: A systematic review. IEEE Reviews in Biomedical Engineering, 10:3122522, 2021.

[40] Antonio Ortega, Pascal Frossard, Jelena Kovacevic, José M F Moura, and Pierre Vandergheynst. Graph signal processing: Overview, challenges, and applications. Proceedings of the IEEE, 106(5):808–828, 2018.

[41] Xiaowen Dong, Dorina Thanou, Laura Toni, Michael Bronstein, and Pascal Frossard. Graph signal processing for machine learning: A review and new perspectives. IEEE Signal Processing Magazine, 2020.

[42] Stefan Bloemheuvel, Jurgen van den Hoogen, and Martin Atzmueller. A computational framework for modeling complex sensor network data using graph signal processing and graph neural networks in structural health monitoring. Applied Network Science, 6(97), 2021.

[43] Thomas N Kipf and Max Welling. Semi-supervised classification with graph convolutional networks. In International Conference on Learning Representations, 2017.

[44] Michael Defferrard, Xavier Bresson, and Pierre Vandergheynst. Convolutional neural networks on graphs with fast localized spectral filtering. In Advances in neural information processing systems, pages 3844–3852, 2016.

[45] Mordjane Boukhet. Discrimination and Sequencing of Polymers with Biological Nanopores. Doctoral dissertation, Albert-Ludwig-Universität Freiburg Germany and Université Cergy-Pontoise France, 2018.

[46] Ángel Díaz Carral, Martin Roitegui, Ayberk Koc, Mag-nus Ostertag, and Maria Fyta. Concurrent analysis of electronic and ionic nanopore signals: blockade mean and height. Nano Express, 5(2):025020, 2024.

